# Role of small intronic RNAs in the crosstalk between immune cells and β-cells during type 1 diabetes development

**DOI:** 10.64898/2025.12.19.695414

**Authors:** Shagun Poddar, Flora Brozzi, Cristina Cosentino, Cécile Jacovetti, Claudiane Guay, Jérôme Perrard, Romano Regazzi

## Abstract

Small non-coding RNAs, such as microRNAs and tRNA-derived fragments, are key regulators of cellular processes, but the functions of small intronic RNAs (sinRNAs), a recently identified RNA class, remain largely unknown. Here, we report that two sinRNAs, sinR-D and sinR-T, are upregulated in pancreatic β-cells of NOD mice, a well-established model of type 1 diabetes. Using in vivo RNA-tagging, we demonstrate that these sinRNAs are packaged into extracellular vesicles released by infiltrating CD4⁺ T lymphocytes and subsequently delivered to β-cells during the early stages of autoimmune attack. Functional analyses revealed that overexpression of sinR-T has little effect on β-cell viability, whereas sinR-D markedly increases β-cell apoptosis. This finding suggests that the transfer of sinR-D contributes to β-cell destruction and the onset of type 1 diabetes. Furthermore, pull-down experiments with biotinylated sinRNAs identified Ago2, a core component of the RNA-induced silencing complex (RISC), as a binding partner of sinR-D, indicating mechanistic parallels with microRNA-mediated regulation. Collectively, our data uncover a novel role for sinRNAs as extracellularly transferred regulators of β-cell fate, expanding the repertoire of small RNAs implicated in the initiation of type 1 diabetes.

## 1. INTRODUCTION

Type 1 diabetes (T1D) is an autoimmune disorder characterized by progressive loss of pancreatic β-cells, resulting in insulin deficiency and chronically elevated blood glucose levels [1, 2]. The autoimmune attack is initiated by the infiltration of the islets of Langerhans by immune cells that release inflammatory mediators causing dysfunction and apoptosis of β-cells [3]. A better understanding of these events is necessary to permit the development of novel strategies to prevent and treat T1D.

Different classes of non-coding RNAs (ncRNAs), including microRNA (miRNAs), tRNA-derived fragments (tRFs), long non-coding RNAs and circular RNAs, contribute to the control of β-cell function and have been proposed to be involved in the development of various forms of diabetes [4–8]. Indeed, the level of several ncRNAs is altered in the islets of prediabetic NOD mice, a well characterized model of T1D [9–11]. Many of these changes are triggered by chronic exposure of β-cells to proinflammatory cytokines, such as IL-1β, TNFα and IFNγ [12–15]. In addition, we found that a group of miRNAs produced by CD4^+^ T lymphocytes invading the islets can be directly transferred to β-cells via extracellular vesicles (EVs) [16]. Blockade of these miRNAs in the receiving β-cells permitted to partially prevent T1D occurrence in NOD mice, suggesting that this mechanism contributes to the development of the disease. Using an RNA-tagging approach, we recently reported that beside miRNAs, different tRFs are also shuttled from immune cells to β-cells, promoting apoptosis of insulin-secreting cells [17].

In addition to tRFs, this approach led to the identification of other small RNAs shuttling from T lymphocytes to β-cells. These include a group of small intron-derived RNAs (sinRNAs), a newly discovered class of non-coding RNAs that is still poorly characterized [18]. The purpose of the present study was to determine the functional role of these small ncRNAs in β-cells and to elucidate their mode of action. Our data further expand the repertoire of small non-coding RNAs regulating the activity of insulin-secreting cells and suggest a possible contribution of sinRNAs in T1D diabetes pathogenesis.

## 2. MATERIALS AND METHODS

### Animals

ARRIVE guidelines were followed. Male C57BL/6NRj mice (aged 12 – 14 weeks) and female NOD.Cg-Prkdc scid/Rj (aged 4 weeks and 8 weeks) were purchased from Janvier and were housed on a 12-h light/dark cycle under standard conditions with *ad-libitum* chow diet and water access. All procedures were performed in agreement with the NIH guidelines and according to the Swiss national legislation and they were approved by the Swiss federal food safety and veterinary offices (animal license numbers VD2744×3 and VD2495×4). Animal euthanasia has been performed by pentobarbital injection in accordance with the Swiss veterinary guidelines.

### Cell lines

The murine MIN6B1 cell line was maintained in DMEM-GlutaMAX containing 25 mM glucose and 4 mM L-glutamine medium supplemented with 15% FBS, 70 µM β-mercaptoethanol, 100 U/mL penicillin and 100 μg/mL streptomycin. MIN6B1 cells were cultured in humidified 5% (vol/vol) CO_2_, 95% (vol/vol) air at 37°C and tested negative for mycoplasma contamination.

### T Lymphocyte isolation and cell culture

Mouse CD4^+^/CD25^-^ T-cells were purified from spleen of C57BL/6NRj mice using CD4^+^/CD25^-^Regulatory T Cell Isolation Kit (Miltenyi Biotec, Germany). CD4^+^/CD25^–^ T cells were cultured in RPMI 1640 medium containing 10% EV-depleted FBS, 100 µg/mL streptomycin and 100 IU/mL penicillin, and stimulated with 20 ng/mL IL-12, 200 IU/mL IL-2, 2 μg/ml anti-CD28 and 5 μg/ml anti-CD3. Media and cells were harvested after 3 and 7 days for EV isolation and/or RNA extraction.

### Isolation of pancreatic islets and β-cell sorting

Mouse islets were isolated as previously described [19] by digesting the pancreas with collagenase, followed by separation using Histopaque density gradient and handpicking. Islet cells were dispersed by incubation with Ca^2+^/Mg^2+^ free phosphate buffered saline, 3 mM EGTA and 0.025% trypsin for 2-3 min at 37⁰C and were then cultured in RPMI 1640 GlutaMAX medium supplemented with 10% FBS, 1 mM sodium pyruvate, 100 U/mL penicillin and 100 μg/mL streptomycin, and 10 mM Hepes, as previously described [16, 17]. Sample preparations from NOD/ShiLtJ (aged 4 and 8 weeks, Janvier) and FAC (Fluorescence- Activated Cell) – sorted β-cells have been previously described [17].

### Isolation of extracellular vesicles

EVs were isolated using a method described previously [16, 17]. Briefly, the culture media (containing EV-depleted FBS) from mouse T cells were first centrifuged at 300 x g for 6 minutes, and then at 2,000 x g for 10 minutes. The supernatants were then centrifuged at 10,000 x g for 30 minutes to eliminate cell debris, and subsequently at 100,000 x g for 2 hours. The resulting pellets containing the EVs were washed twice with PBS and subjected to a final centrifugation at 100,000 x g for 2 hours.

### Cell transfection and treatment

Dispersed mouse islet cells were transfected with scrambled or custom-designed RNA oligonucleotides (IDT) mimicking the sequence of sinR-D2/D3 or sinR-T using Lipofectamine 2000 and incubated for 48h before RNA extraction or functional assays. Oligonucleotide sequences are provided in Supplementary Table 2. The effect of pro-inflammatory cytokines was examined by incubating the cells with IL-1β (0.1 ng/mL), TNF-α (10 ng/mL), and IFN-γ (30 ng/mL) for 24h. For the treatment with EVs, islet cells were exposed to EVs (1×10^11^/mL) for 24h.

### Immunofluorescence

The cells were cultured in removable 12-well chamber slides (Cat # 81201, Ibidi, Germany) coated with Poly-L-Lysine and laminin, and were fixed with methanol. They were then permeabilized using saponin and double-stained with 1:200 mouse anti-insulin (cat # 66198-1, Proteintech, USA) and 1:500 rabbit anti-cleaved-caspase 3 (cat #9661S, Cell Signaling Technology, USA) antibodies. AlexaFluor 488 anti-mouse and AlexaFluor 555 anti-rabbit (1:400, Invitrogen, USA) were used as secondary antibodies. Cell nuclei were stained with Hoechst 33342 (1 μg/ml, Invitrogen). Coverslips were mounted on microscope glass slides with Fluor-Save mounting medium (VWR International SA) The images were acquired with a Zeiss Axiovision fluorescence microscope.

### Insulin secretion and insulin ELISA

Transfected mice islet cells and MIN6 cells were first incubated at 37 °C for 60 min in a Krebs–Ringer bicarbonate buffer (KRBH) containing 25 mM HEPES, pH 7.4, 0.1 % BSA, and 2 mM glucose. Thereafter, the cells were incubated at 37 °C for 45 min in KRBH-BSA solutions with 2 mM (basal) or 20 mM (stimulatory) glucose. After incubation, supernatants were collected. The cells kept at basal glucose were harvested using acid ethanol (75 % ethanol, 0.55 % HCl), and those incubated at stimulatory conditions were lysed using Triton X-100 lysis buffer to determine insulin and protein contents, respectively. Insulin levels were measured by ELISA according to the manufacturer’s protocol (Mercodia, Sweden) and cellular protein contents by Bradford assay.

### Pull-down assay and western blotting

To elucidate the interaction between sinRNAs and Ago2, a pull-down assay was performed using 5’- biotinylated oligonucleotides mimicking the sequence of sinR-D or sinR-T or their scrambled controls as previously described [20]. Briefly, 1 mg of MIN6B1 cell lysate was prepared using lysis buffer (50 mM Tris–HCl pH 7.5, 150 mM NaCl, 1% NP40, 0.1% SDS, 0.5% sodium deoxycholate, 1 mM DTT, protease inhibitor cocktail, 80 U/ml RNase inhibitor) and incubated with 3 µg of oligonucleotides for 1 h at room temperature on a vertical rotator. Further, 50 μL of streptavidin dynabeads (M-280 Streptavidin, 11206D, Invitrogen) was added to the samples and incubated again for 1h at room temperature on a vertical rotator. The beads were thoroughly washed with a solution containing 50 mM Tris-HCl (pH 7.5), 150 mM NaCl, 1% NP-40, 0.1% SDS, and 0.5% sodium deoxycholate. Subsequently, the proteins were eluted by heating the samples in Laemmli buffer at 95°C for 5 minutes. Protein extracts were loaded on SDS-PAGE and transferred to nitrocellulose membranes (BioRad). The membranes were blocked using 5% BSA in TBST buffer and then incubated with rabbit anti-Ago2 antibody (cat# ab186733, Abcam, UK), overnight at 4°C followed by incubation for 1h at RT with the secondary antibody AlexaFluor 555 anti-rabbit (1:400, Invitrogen, USA). The blots were developed using ECL substrate (cat# 34075, Thermo Fisher, USA).

### Small RNA seq bioinformatics analysis

Sequencing data was filtered to retain reads between 16 and 55 nucleotides in length as previously described [17]. Sequences were further filtered to include only those with a minimum of 5 counts in at least 4 samples, yielding 63,367 unique sequences. To focus on mouse-derived fragments and exclude contaminants or sequences of unknown origin, only reads matching the reference *Mus musculus* genome (mm10) were retained. Differential expression analysis was conducted using DESeq2 on three sets of sequences: all 7,900 fragments, 242 tRNA-derived fragments, and 217 microRNA-derived fragments [17] with normalization validated through miR-238 C. elegans spike-in control. Statistical significance was determined using an adjusted p-value threshold of ≤ 0.01, identifying differentially abundant fragments in EU-tagged RNA extracted from receiving beta cells.

### Proteomics analyses

All raw MS data together with raw output tables are available via the Proteomexchange data repository (www.proteomexchange.org) with the accession PXD065257 (reviewers can access the data at: https://www.ebi.ac.uk/pride/login; Username; Password: FdtVmQYPGyRV). Samples were digested following the SP3 method [21] using magnetic Sera-Mag Speedbeads (Cytiva 45152105050250, 50 mg/ml). Briefly, samples were diluted with SP3 buffer (2% SDS, 10mM DTT, 50 mM Tris, pH 7.5) and heated 10 min at 75°C. Proteins were then alkylated with 30mM iodoacetamide for 45 min at RT in the dark. Proteins were precipitated on beads with ethanol (final concentration: 60 %). After 3 washes with 80% ethanol, beads were digested in 50ul of 100 mM ammonium bicarbonate with 1.0 ug of trypsin (Promega #V5113). After 1h of incubation at 37°C, the same amount of trypsin was added to the samples for an additional 1h of incubation. Supernatants were then recovered and two sample volumes of isopropanol containing 1% TFA were added to the digests. The samples were then desalted on a strong cation exchange (SCX) plate (Oasis MCX; Waters Corp., Milford, MA) by centrifugation to remove traces of SDS. After washing with isopropanol/1%TFA and 2% acetonitrile/0.1% FA, peptides were eluted in 200ul of 40% MeCN, 59% water, 1% (v/v) ammonia, and dried by centrifugal evaporation. LC-MS/MS analyses were carried out on a TIMS-TOF Pro (Bruker, Bremen, Germany) mass spectrometer interfaced through a nanospray ion source to an EvoSep One liquid chromatography system (EvoSep, Odense, Denmark). Peptides were separated on a reversed-phase Aurora Elite C18 column (15 cm, 75 μm ID, 1.7um, IonOpticks) at a flow rate of 200 nl/min. Data-independent acquisition was carried out using a method similar to the DIA-PASEF method reported previously [22]. Identification of peptides directly from DIA data was performed with Spectronaut 19.9 with the Pulsar engine using the “deep” setting and searching the reference mouse proteome (www.uniprot.org) database of February 6^th^, 2025 (54’739 sequences), and a contaminant database containing the most usual environmental contaminants and enzymes used for digestion [23]. For identification, peptides of 7-52 AA length were considered, cleaved with trypsin/P specificity and a maximum of 2 missed cleavages. Carbamidomethylation of cysteine, methionine oxidation and N-terminal protein acetylation were the modifications applied. Ion mobility for peptides was predicted using a deep neural network and used in scoring.

Peptide-centric analysis of DIA data was done with Spectronaut 19.9 using the library generated by Pulsar from DIA data. Peptide quantitation was based on XIC area, for which a minimum of 1 and a maximum of 3 precursors were considered for each peptide, from which the mean value was selected. Quantities for protein groups were derived from inter-run peptide ratios based on MaxLFQ algorithm [24]. Global normalization of runs/samples was done based on the median of peptides. All subsequent analyses were done with an in house developed software tool (https://github.com/UNIL-PAF/taram-backend). Contaminant proteins were removed, and quantity values for protein groups generated by Spectronaut were log2-transformed. Missing values were imputed based on a normal distribution with a width of 0.3 standard deviations (SD), down-shifted by 1.8 SD relative to the median.

### Statistical analysis

Data are presented as means ± SD. A two-tailed one-sample t-test was conducted to compare the datasets against a control value set at 1. Unpaired t-tests were used for comparing two datasets. One-way Anova followed by Dunnett post hoc test was used for multiple dataset comparison. Differences between datasets were considered statistically significant if the p-value was less than 0.05.

## 3. RESULTS

NOD mice are a well-known T1D model characterized by progressive infiltration of immune cells in the islets of Langerhans, causing β-cell loss and culminating in diabetes appearance [9–11, 25]. We recently reported that during the initial phases of T1D, different tRFs are transferred from immune cells to insulin-secreting cells, potentially contributing to β-cell elimination and to the development of the disease [17]. To identify the RNAs transferred to β-cells during the autoimmune attack, CD4^+^ T-cells isolated from NOD BDC2.5 mice were incubated with 5’-ethynyl uridine (EU), a nucleotide derivative incorporated in RNAs during transcription. Upon inoculation into NOD.SCID mice, CD4^+^ T-cells from NOD BDC2.5 mice trigger the appearance of autoimmune diabetes [26]. Two days after the adoptive transfer, when the receiving mice were still in the initial phases of the disease, β-cells were isolated by FACS and EU-tagged RNA molecules shuttled from CD4^+^ T cell to β-cells were purified on streptavidin beads. A schematic representation of the experiment can be found in Fig.1A. In addition to the previously studied tRFs [17], bioinformatic analysis of the RNAs shuttling from T lymphocytes to β-cells revealed an enrichment of small intron-derived RNAs (sinRNAs) (Supplementary table 1; Fig.1B). The three RNAs most significantly transferred to β-cells listed as sinR-D1, sinR-D2 and sinR-D3 map to an intronic region of the *Dnaaf1* gene (Fig. S1A). This sequence is part of an active enhancer validated by STARR-seq (Self-Transcribing Active Regulatory Region sequencing) [27]. Sequence alignment revealed that sinR-D1, D2, and D3 share a conserved 16-nt core sequence (CTCGGTAGAACCTCCA), with differences restricted to one or two nucleotides at the 5′ end. Given this minimal variability and their origin from the same intronic region of the *Dnaaf1* gene, we inferred a shared biogenesis. Therefore, for downstream detection and quantification assays, we selected the sinR-D3-specific primer, as it provides broad coverage of this sinRNA family while avoiding redundancy in primer design. Another interesting candidate is sinR-T which is 24 nt in length and maps to the intronic region of the *Ttll11* gene (Fig. S1B). These sinRNAs are detectable in various tissues (Fig.1C), suggesting that their expression is not restricted to immune cells and β-cells. To verify the results obtained by small RNA-seq, we performed qRT-PCR analysis on the EU-tagged RNAs released by CD4^+^ T lymphocytes and recovered in β-cells two days after the adoptive transfer of the immune cells. The results obtained confirmed the finding of the RNA-seq (Fig. S2)

**Fig. 1.**
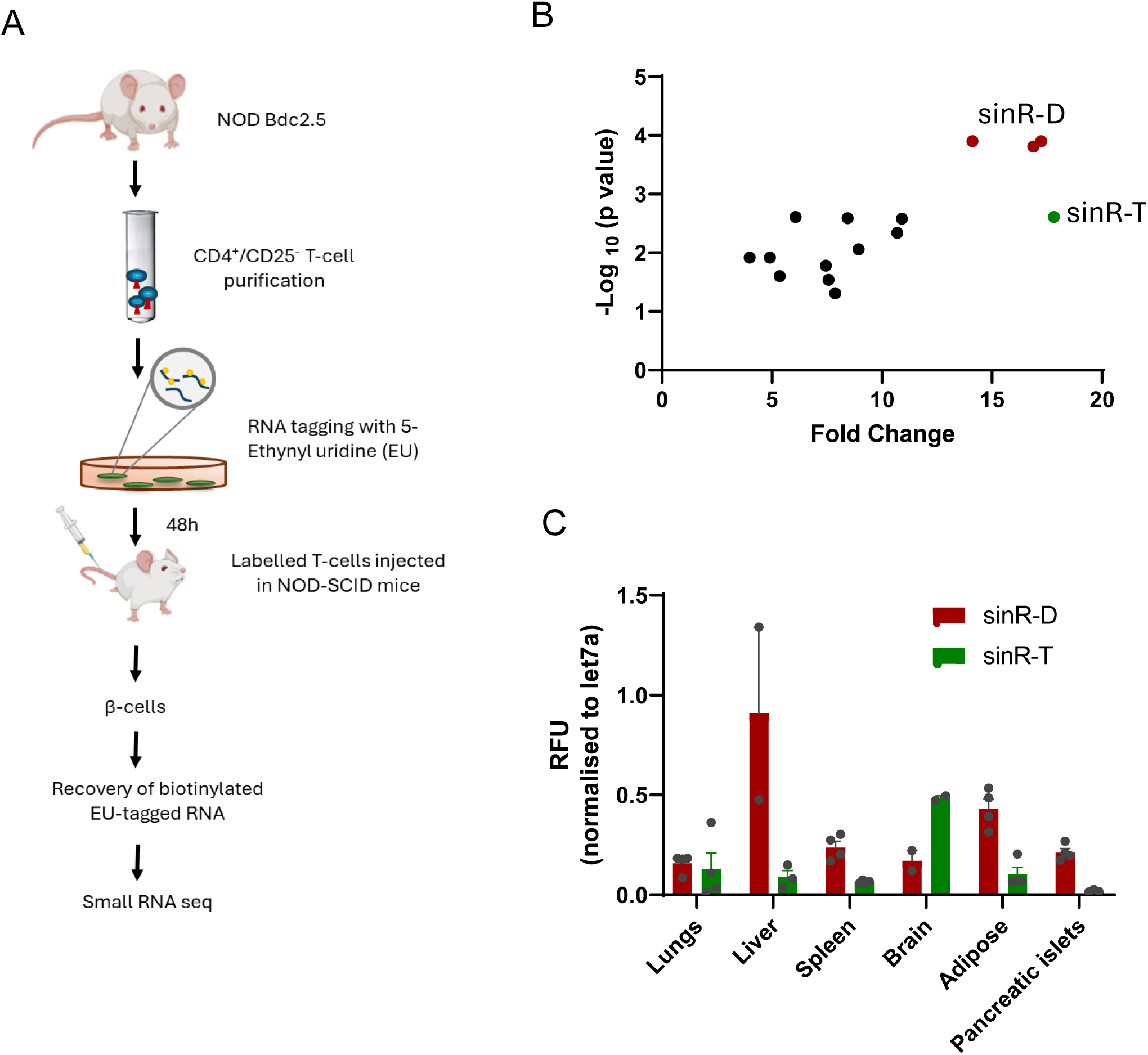
Identification of small RNAs transferred *in vivo* from CD4^+^/CD25^-^ T cells to pancreatic β-cells. (A) Schematic representation of *in vivo* T cell adoptive transfer and identification of EU-tagged RNA in pancreatic β-cells. As previously described [17], CD4^+^/CD25^-^ T cells were purified from NOD.Cg-Tg (TcraBDC2.5, TcrbBDC2.5) mice and were incubated with 200 μM ethynyl uridine (EU) for 48h. The labelled T cells were injected in the tail vein of NOD.CB-17-Prkdc.scid/Rj mice. Saline solution was injected intravenously as control. The pancreatic islets were isolated after 48h and β-cells purified FACS. EU-tagged RNAs were biotinylated using a Click-iT Nascent RNA Capture kit and then isolated on streptavidin-coated beads. The eluted RNA was used for library preparation and small RNA sequencing. (B) Plot showing differential expression of small RNA fragments (16–29 nucleotides) in EU-tagged RNA extracted from the receiving β-cells. Each point represents a small RNA fragment, with the x-axis displaying fold changes and the y-axis -log₁₀ adjusted p-values. Red points highlight the three upregulated sequences of sinR-D and the green point sinR-T. As internal control, C. elegans miR-238 RNA sequence was spiked into each sample, and its EU-tagged sequence was used for normalization across samples. (C) To demonstrate widespread expression of the selected sinRNAs, tissue samples were collected from C57BL6 mice and sinRNA levels were measured using qRT-PCR. The relative abundance of sinRNAs in each tissue was normalised to Let7a. The results are expressed as Relative Fluorescent Units (RFU).

As these sinRNAs are transferred from immune cells to β-cells, we examined whether their level is increased in the islets of prediabetic NOD mice (8 weeks old) compared to the islets of young NOD mice (4 weeks old) that are devoid of insulitis [28]. As shown in Fig.2A, the levels of sinR-D3 and sinR-T were increased significantly in the islets of prediabetic NOD mice where immune cells have begun to invade the islets. To rule out age-related effects, we also assessed sinRNA levels in age-matched NOD-SCID mice at 4 and 8 weeks of age, which do not develop diabetes due to the absence of functional immune cells. Notably, no significant changes in sinR-D3 or sinR-T levels were observed in NOD-SCID islets (Fig. S3), suggesting that the upregulation is associated with immune infiltration rather than aging.

**Fig. 2.**
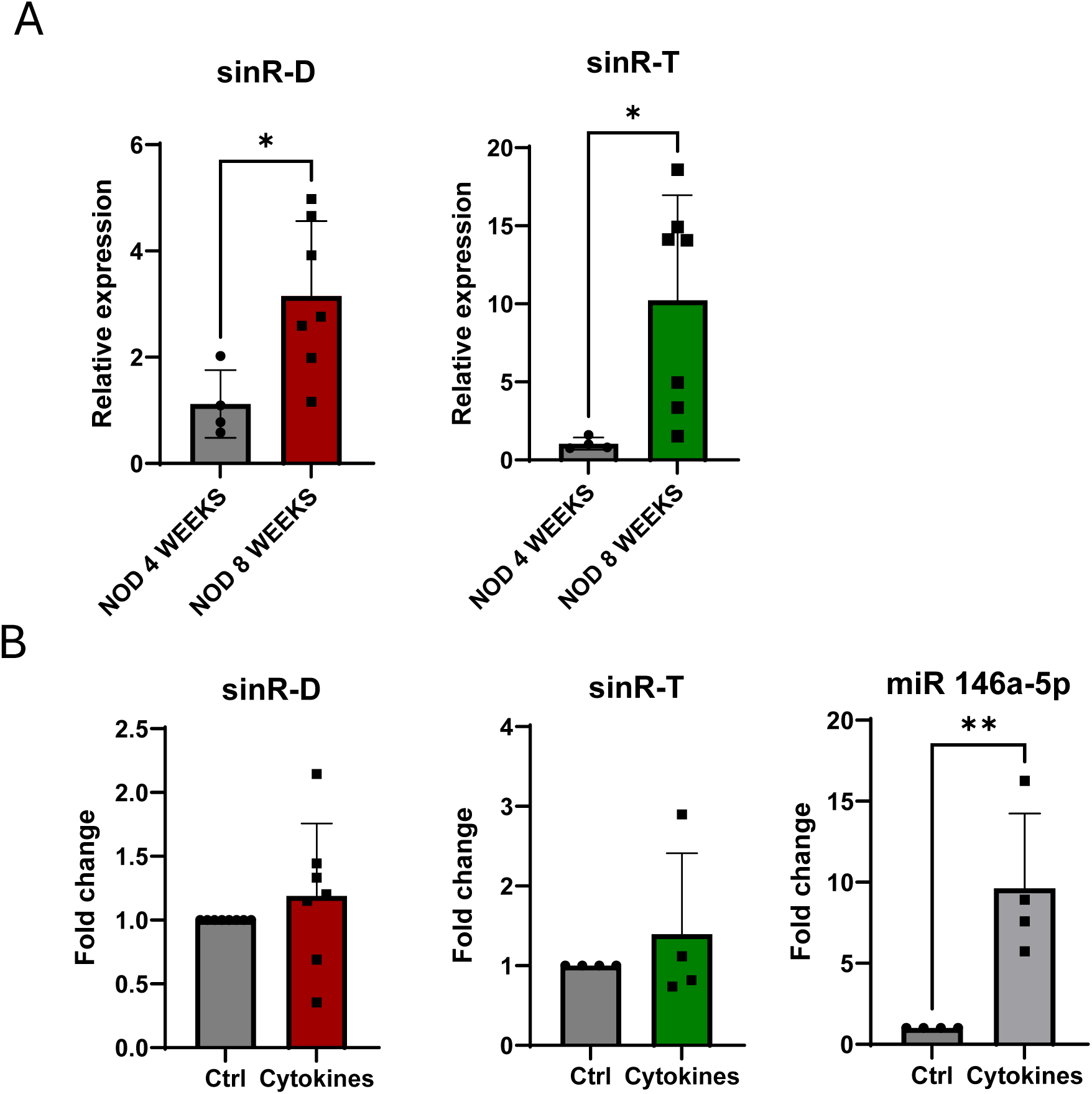
Levels of sinRNAs in prediabetic NOD mice and cytokine-treated islets. (A) sinRNA expression levels were quantified in NOD mice at 4 weeks and 8 weeks by qRT-PCR, with miR-184 serving as the normalization control. Data shown are the means ± SD, n=4-7, *p<0.05. (B) Islet cells of C57BL/6NRj mice incubated with or without cytokines (IL-1β, IFN-γ and TNF-α) for 24h prior to RNA extraction. qRT-PCR was performed to measure sinRNA levels, with Let7a used for normalization (n=4-7 independent experiments) mean ± SD.

Proinflammatory cytokines released by the infiltrating immune cells play a crucial role in regulating the expression of different genes and small ncRNAs in β-cells [16, 17]. To examine this possibility, we checked the effect of proinflammatory cytokines on the levels of these sinRNAs. Pancreatic islets isolated from C57BL6N mice were dissociated and treated with a mix of proinflammatory cytokines (IL-1β, TNF-α and IFN-γ). After 24h incubation, RNA was extracted and the level of the sinRNAs measured by qRT-PCR. We found that, in contrast to the positive control miR-146a, the level of these sinRNAs was not significantly affected by cytokine exposure (Fig.2B), suggesting that the increase observed in β-cells after the adoptive transfer of CD4^+^ T cells and in the islets of prediabetic NOD mice is not caused by the presence of proinflammatory cytokines.

EVs mediate the exchange of signals between different cell types [29]. T cells secrete EVs with distinct cargoes, including miRNAs that contribute to increase the apoptotic rate of recipient islet cells [16]. The expression of sinR-D3 and -T was detected both in the RNA extracted from activated CD4^+^/CD25^-^ T lymphocytes and in the EVs released by them (Fig. 3A). Moreover, the levels of sinRNAs were increased in a time dependent manner upon activation of CD4^+^/CD25^-^T lymphocytes as shown in Fig.3B. To investigate whether the EVs released by T cells could mediate the rise of sinRNAs in islet cells, CD4^+^/CD25^-^ T lymphocytes were extracted from the spleen of C57BL6N mice and were activated with IL2, IL12 and anti-CD3 and anti-CD28 beads. The EVs released from T lymphocytes were then purified by ultra-centrifugation and added to the culture medium of dispersed mouse islet cells. After 24h, RNA was extracted and the level of the sinRNAs was assessed by qRT-PCR. These measurements showed that the level of the sinR-D3 in islet cells is increased in the presence of the EVs released by activated CD4^+^/CD25^-^ T lymphocytes (Fig.3C). Although a similar trend was observed for sinR-T, the effect was smaller and did not reach statistical significance. To evaluate whether this process also occurs in humans, human islet cells were incubated with exosomes (EV-hT) derived from CD4⁺ T cells isolated from the human blood donors, as previously described [16]. Consistent with the findings in mice, exposure of human islet cells to EV-hT facilitated the transfer of sinR-D3, whereas transfer of sinR-T was comparatively limited (Fig.3D). Taken together, these in vitro findings support our previous *in vivo* results demonstrating that sinRNAs are transferred from T lymphocytes to pancreatic islet cells via EVs during the development of T1D.

**Fig. 3.**
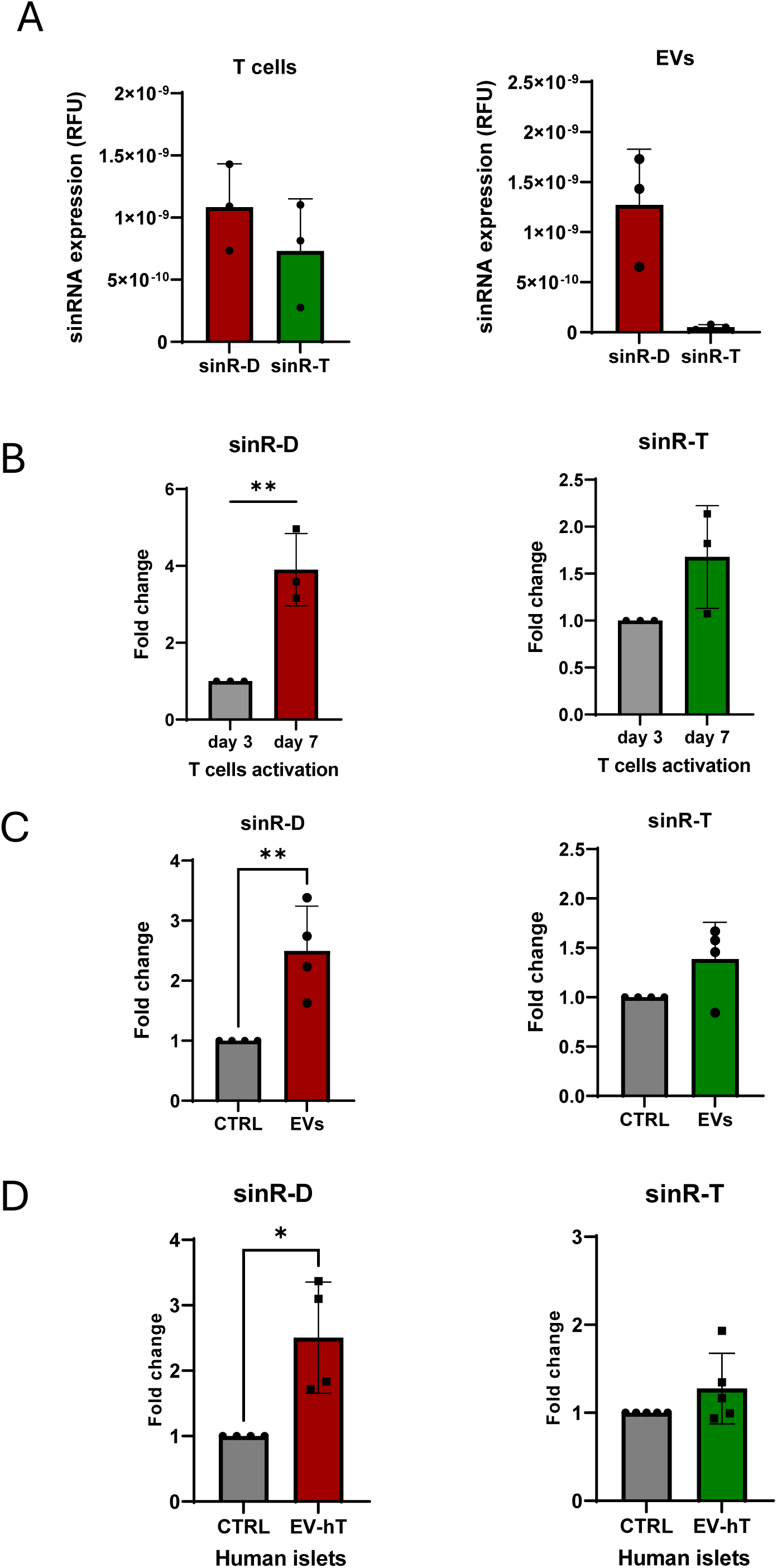
Transfer of sinRNAs from T cells to islet cells via extracellular vesicles. (A) Expression levels of sinRNAs were measured in 100ng RNA extracted from activated CD4^+^/CD25^-^ T lymphocytes and from EVs released by these cells. The values are calculated using 2^^-Ct^ method and presented as Relative Fluorescent Units (RFU). (B) RNA was extracted from CD4^+^/CD25^-^ T lymphocytes following activation of 3 days (3d) and 7 days (7d) and the levels of the sinRNAs measured by qPCR using Let7a for normalisation (n=3). (C-D) qRT-PCR analysis of sinRNA levels in pancreatic islet cells following a 24-hour incubation with extracellular vesicles (EVs) released by CD4⁺ T cells—(C) mouse islets with EVs from murine CD4⁺ T cells and (D) human islets with EVs from CD4⁺ T cells of human donors. Let7a was used normalization. Data are presented as mean ± SD, with n=4 biological replicates. Statistical significance was determined as \*\**p*< 0.01.

As demonstrated in our previous work [16, 17], exposure to T cell-derived EVs leads to β-cell death. We hypothesized that these sinRNAs may contribute to this effect upon their transfer. To investigate the functional relevance and the impact of these sinRNAs, we overexpressed them in mouse islet cells using oligonucleotides mimicking the sequence of sinR-D3 and sinR-T and we then assessed the impact on β-cell death. Immunofluorescence analysis with anti-insulin and anti-cleaved caspase-3 antibodies revealed a significant increase in apoptosis upon sinR-D3 over-expression (Fig.4A). In contrast, no significant difference was observed when the level of sinR-T was increased. These findings suggest that the rise of sinR-D3 resulting from the delivery of T cell EV cargoes may contribute to β-cell death occurring during the initial phases of the disease. Insulin content (Fig.4B) and glucose-induced insulin secretion (Fig.4C) were not affected by overexpression of sinR-D3. Similar findings were obtained by overexpressing sinR-D3 alone or in combination with sinR-D2 in MIN6 cells (Figure S4).

**Fig. 4.**
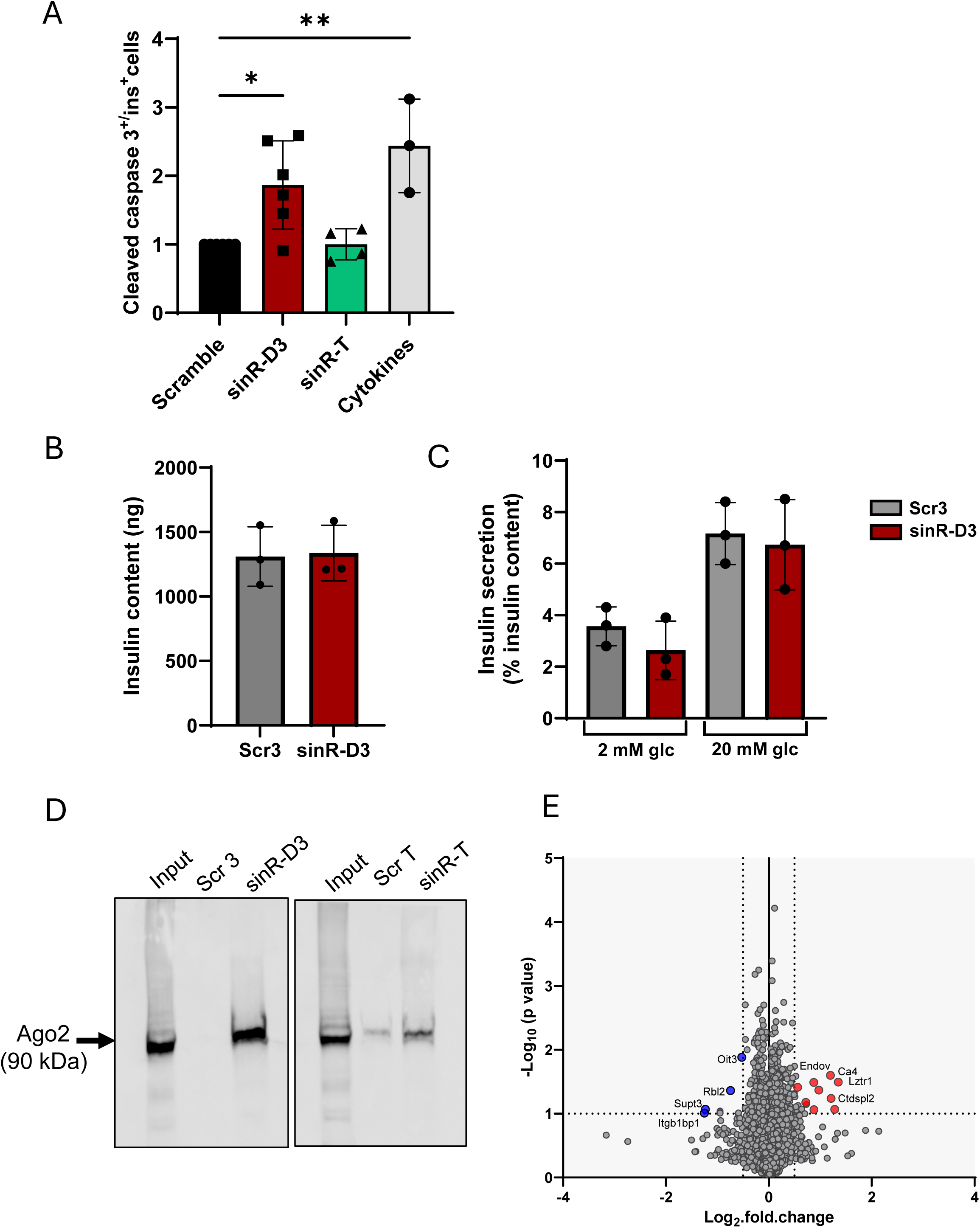
Functional impact and mode of action of sinRNAs. (A) Mouse islet cells were transfected with sinRNA mimics or with a scrambled oligonucleotide sequence and incubated for 48h. Where indicated, islet cells were exposed to pro-inflammatory cytokines (IL-1β, IFN-γ, and TNF-α) for 24h. At the end of the incubation, the coverslips were stained with anti-insulin and anti-cleaved caspase-3 antibodies, and the percentage of cleaved caspase-3 positive β-cells was determined. Fold changes were then calculated relative to the scrambled condition. *p<0.05, n=3-6, mean ± SD. (B) Insulin content of mouse islets transfected sinR-D3 or with a scrambled oligonucleotide sequence (Scr3). (C) Glucose-induced insulin release of transfected mouse islet cells (as mentioned above) in response to 2 mM glucose (basal) or to 20 mM glucose (stimulatory). (D) Western blot analysis demonstrating the enrichment of Ago2 protein in MIN6 cells incubated with either biotinylated sinRNA mimics or the corresponding scrambled control sequences, followed by pull-down with streptavidin beads. Input lane represents 10% of the total cell lysate. Data shown is a representative image from three independent experimental replicates. (E) Volcano plot representing differentially expressed proteins identified by proteomic analysis of pancreatic islets overexpressing both sinR-D2 and sinR-D3 compared to control cells transfected with length-matched scrambled sequences. Significantly upregulated proteins are shown in red, and significantly downregulated proteins are shown in blue (p < 0.05).

Although the exact mechanism of action of sinRNAs remains largely unexplored, few reports suggest that some of them could bind Ago2 and act as miRNAs [30] and other small RNA classes [31–33]. Indeed, evidence for the physical association of some sinRNAs with Ago2 and for their capacity to regulate gene expression has been recently reported [18]. Notably, we found exact sequence matches for sinR-D2 and sinR-D3 in publicly available datasets from the same study [18], specifically in Supplementary Data 7 (Top 5000 15–30 nt Ago2-binding sRNA-OHs identified by TANT-seq in Hepa 1-6 cells) and Supplementary Data 9 (Top 5000 15–30 nt Ago2-binding sRNA-OHs identified by TANT-seq in human 293T cells). These findings support the incorporation of sinR-D2 and sinR-D3 into Argonaute complexes and suggest a conserved potential for functional activity across species and cell types. To verify whether sinR-D3 and sinR-T are able to bind to Ago2, we incubated biotinylated mimics of each sinRNA with lysates of the insulin-producing MIN6B1 cells (Figure S5). We then used streptavidin beads to isolate from the extract the proteins associated to these small RNAs. Western blot analysis using an antibody against Ago2 demonstrated a clear interaction between Ago2 and sinR-D3, while little or no specific interaction was observed with sinR-T (Fig.4D).

Given the observed physical association of sinR-D3 with Ago2, we next aimed to assess the impact of this sinRNA on protein expression. To capture the full range of potential regulatory effects, we conducted overexpression experiments using a combined mixture of sinR-D2 and sinR-D3, which differ by a single nucleotide at the 5′ end. This choice was based on the principle that, similar to miRNAs, small RNA function can be highly sensitive to sequence variation, particularly within the seed region, where even a single nucleotide difference may alter target specificity [34–37]. By including both variants, we aimed to ensure comprehensive representation of potential target interactions. The resulting proteomic changes were then analyzed by quantitative mass spectrometry. Proteomic analysis of sinRNA-overexpressing pancreatic islets revealed a downregulation of RBL2, OIT3, ITGB1BP1, and SUPT3, which are associated with pathways involved in cell cycle regulation, adhesion, and transcription [38–43]—processes whose disruption is known to promote apoptosis (Fig.4E, Supplementary Table 3). Conversely, CA4, LZTR1, ENDOV, CTDSPL2, and ST3GAL5 were significantly upregulated and are linked to stress response, DNA damage, and apoptotic signaling [44–48], suggesting activation of pro-apoptotic mechanisms in response to sinRNA overexpression.

Collectively, these data suggest that, in addition to miRNAs and tRFs, distinct sinRNAs are also transferred from immune cells to pancreatic β-cells during the early phases of T1D. Among them, sinR-D3 was found to interact with Ago2 and may lead to gene regulation that contributes to β-cell death.

## 4. DISCUSSION

The discovery of small ncRNAs has rapidly emerged as a transformative area in molecular biology. These small ncRNAs typically 15 to 50 nucleotides in length, include well-characterized species such as miRNAs. Recent advancements in high-throughput sequencing technologies and bioinformatics tools have expanded the repertoire of identified small ncRNAs across diverse species [18, 49, 50]. This progress has sparked growing interest in newer classes of small ncRNAs derived from longer, structured RNA molecules such as tRFs (derived from transfer RNAs)[51, 52], rRFs (derived from ribosomal RNAs)[53], Y RNA fragments [54], small nuclear RNAs (snRNAs)[55], small nucleolar RNAs (snoRNAs)[56], vault RNAs (vtRNAs)[57], and other less-characterised small ncRNAs [18, 58]. Understanding the functional roles of these emerging small ncRNAs is crucial to elucidating their involvement in physiological and pathological processes.

In this study, we investigated a subset of small ncRNAs derived from the cleavage of intronic sequences, focusing on their role in the early stages of T1D development. Many small ncRNAs are incorporated into EVs and are selectively released into the extracellular space, where they play an important role in intercellular communication [16, 17, 59, 60]. Indeed, the delivery of small ncRNAs carried by EVs has been shown to trigger functional changes in the receiving cells and to contribute to many pathological processes, including the development of different forms of diabetes [61].

Recent evidence indicates that lymphocytes infiltrating the islets of Langerhans during T1D development release EVs containing a distinct set of miRNAs. These miRNAs are transferred to β-cells, triggering the activation of apoptotic pathways [16, 62, 63]. In addition to miRNAs, we recently demonstrated that EVs from CD4^+^ T cells invading the islets also transfer tRFs to β-cells [17]. This suggests that EVs molecular cargo extends beyond the known small ncRNAs, and RNA labelling methodologies are able to unveil unexplored molecules that contribute to cell-to-cell communication.

Our study focused on the functional role of two small RNAs, which are derived from intronic regions and are shuttled between CD4^+^ T cells and β-cells during the initial stages of T1D. Specifically, sinR-D3 originates from the host gene Dnaaf1, and sinR-T is derived from Ttll1. We found that these sinRNAs are expressed in various tissues, with elevated levels in pancreatic islets of prediabetic NOD mice—a well-established model of T1D. Notably, high-throughput sequencing studies have identified sequences corresponding to our sinRNAs in mouse and human cell lines [18], further validating their existence as an underappreciated class of small ncRNAs.

Functionally, we observed that overexpression of sinR-D3 induces apoptosis in pancreatic β-cells, suggesting a regulatory role in insulin-secreting cells and its potential contribution to T1D progression through β-cell death. While the precise molecular mechanisms remain to be fully elucidated, our evidence indicates that sinR-D3 may act similarly to miRNAs, mediating post-transcriptional gene repression. Indeed, pull-down experiments with biotinylated oligonucleotides revealed that sinR-D3 interacts with Ago2, a key component of the RISC complex. These findings align with the observations of Lai et al.[18], who demonstrated that many sinRNAs, including sinR-D, co-immunoprecipitate with Ago2. Although overexpression of sinR-D resulted in changes in the level of proteins potentially involved in apoptosis, the precise mechanisms through which this small ncRNA affects β-cell survival remains to be fully elucidated and would need to be addressed in the future.

Our study expands the repertoire of ncRNAs involved in the crosstalk between immune cells and β-cells, highlighting their regulatory roles in insulin-secreting cells under diabetic conditions. These findings underscore the need to investigate sinRNAs further to understand their contributions to cellular signalling pathways. A deeper understanding of this overlooked class of small ncRNAs could provide significant insights into the pathogenesis of diabetes and other human diseases.

## AUTHORS CONTRIBUTION

SP, FB, CC, CJ, CG and JP contributed to data acquisition, analysis and interpretation. They critically revised the manuscript and agreed to the submission of the article. RR conceived the study, interpreted the data and wrote the manuscript.

## ACKNOWLEDGMENTS

We are grateful to Dr. Nicolas Guex and Dr. Christian Iseli from the Bioinformatics Competence Center at the University of Lausanne, for analyzing the small RNA-sequencing data. We thank Dr. Manfredo Quadroni and the entire team of the Protein Analysis Facility, Faculty of Biology and Medicine, University of Lausanne, Switzerland for excellent expertise, advice, and analysis of the proteomics.

## DATA AVAILABILITY

All the data are available upon request.

---

**Supplementary Figure S1.**
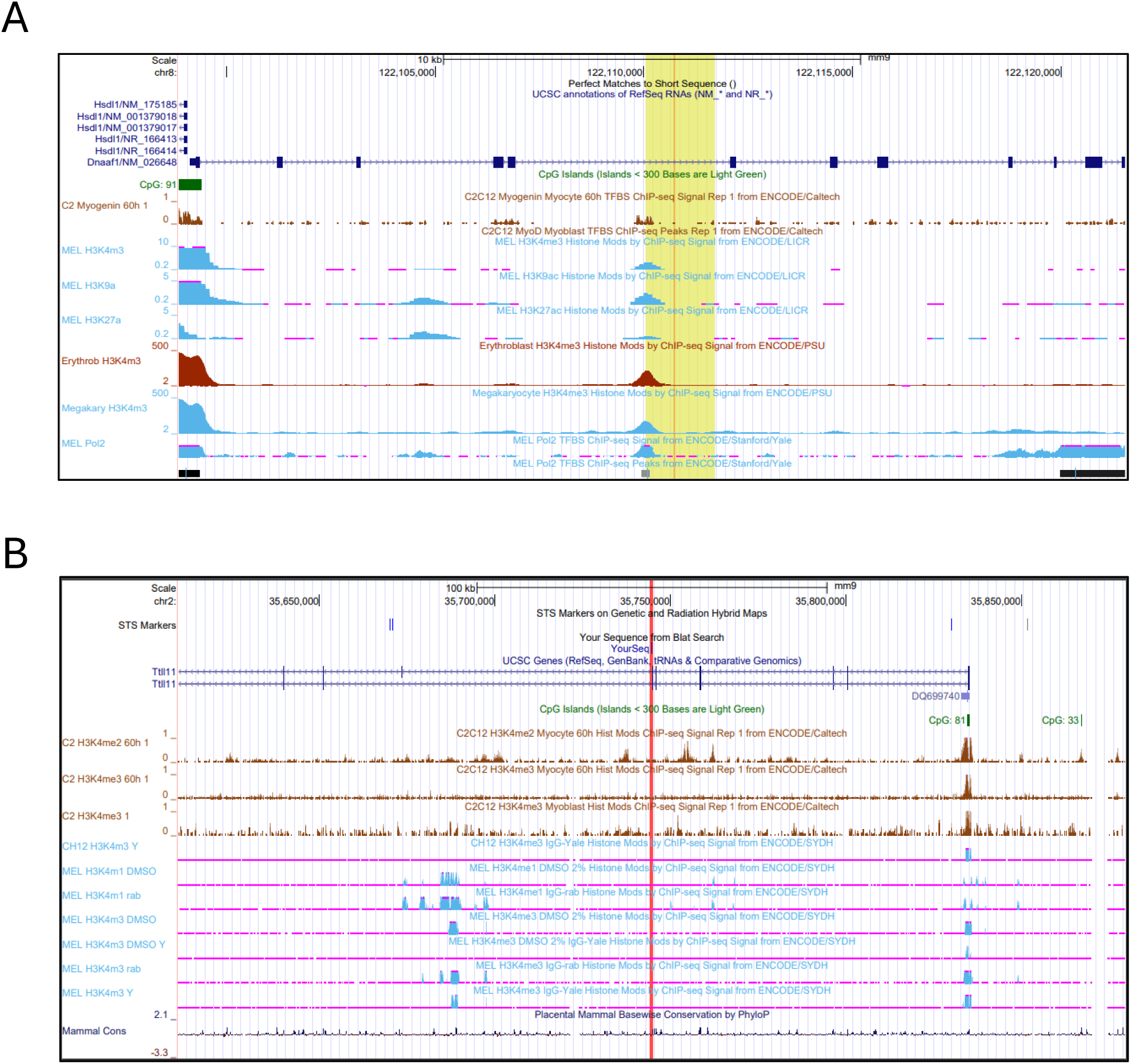
Visualization of sinRNA locations using the UCSC Genome Browser. (a) sinR-D is mapped to intron 5 of the *dnaaf1* gene, indicated by an orange vertical line. The yellow segment highlights STARR-seq peaks, suggesting histone modifications at the start, indicative of a potential regulatory region. (b) sinR-T is located within an intron of the *TTLL11* gene, shown by a red vertical line, with histone modification peaks observed within the intronic region."

**Supplementary Figure S2.**
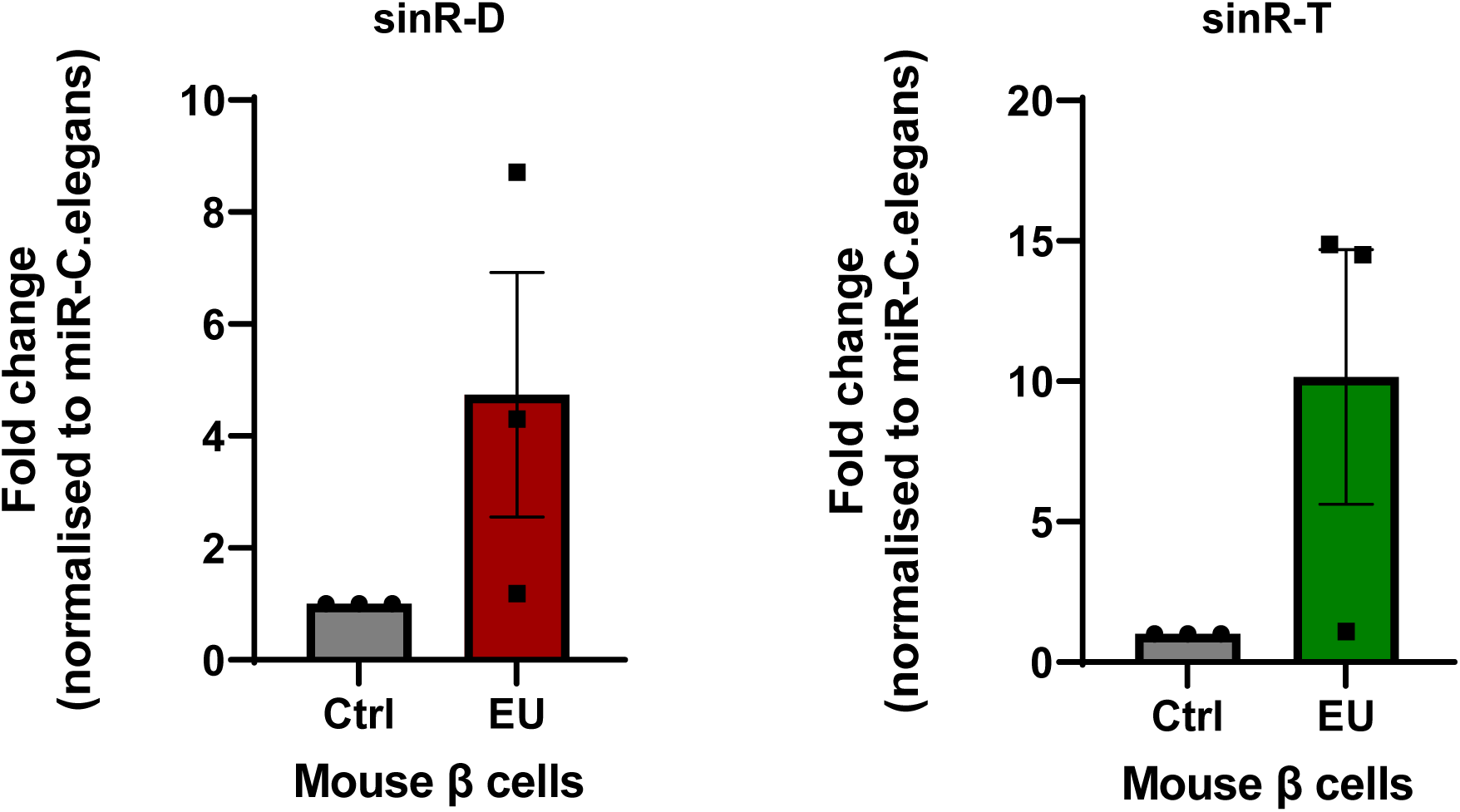
qRT-PCR validation of small RNA-seq results. EU-tagged RNAs from CD4^+^ T lymphocytes were analyzed in beta cells three days post-adoptive transfer of immune cells. The qRT-PCR findings corroborate the RNA-seq data. The sequence of spike-in *C.elegans* miR-238 was used as normalisation control. Data are presented as mean ± SD, n=3.

**Supplementary Figure S3.**
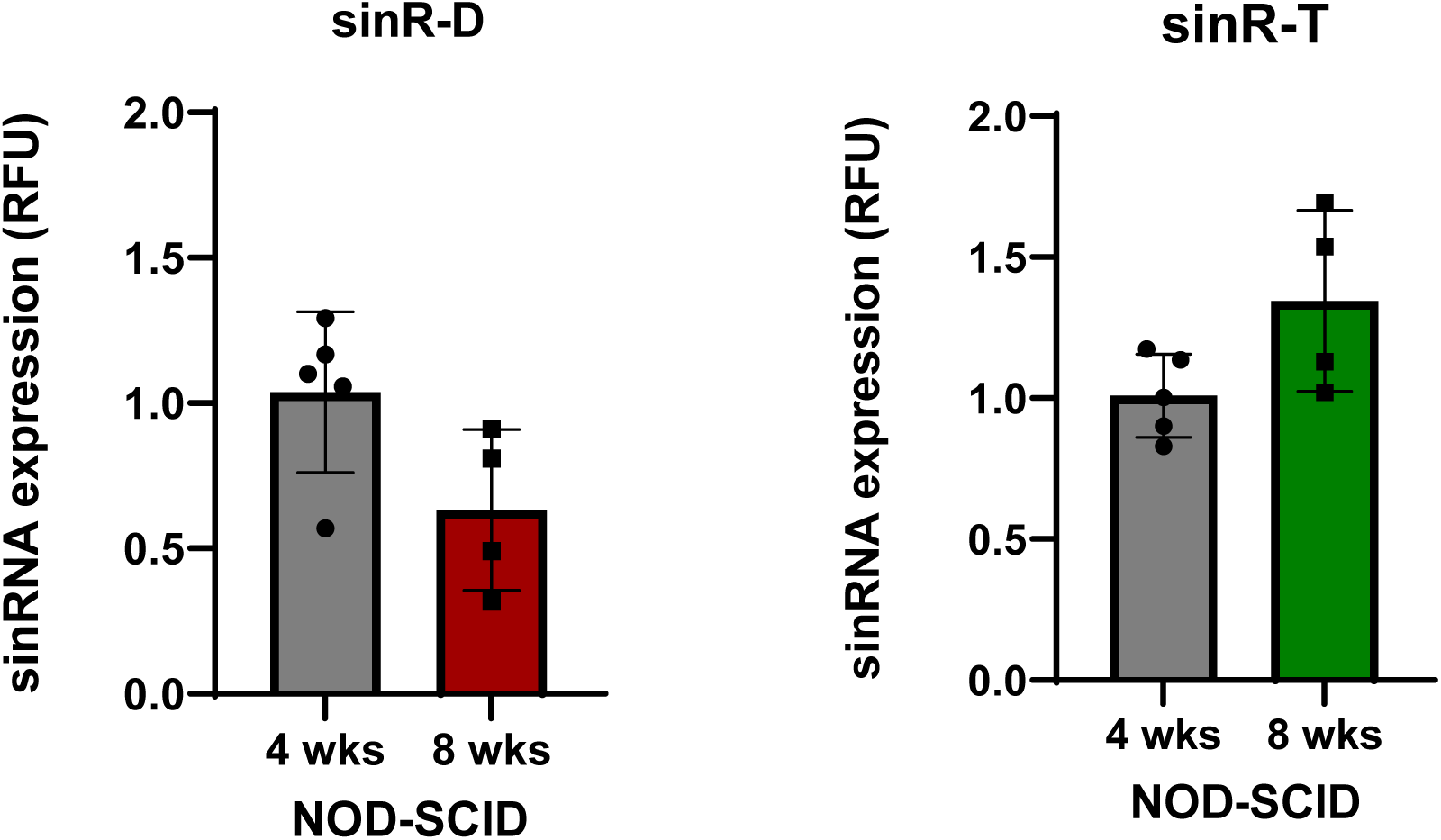
Levels of the selected sinRNAs in NOD-SCID mice islets. sinRNA levels were quantified in NOD-SCID mice at 4 weeks and 8 weeks by qRT-PCR, Let7a was used for normalization, n=4-5, mean ± SD.

**Supplementary Figure S4.**
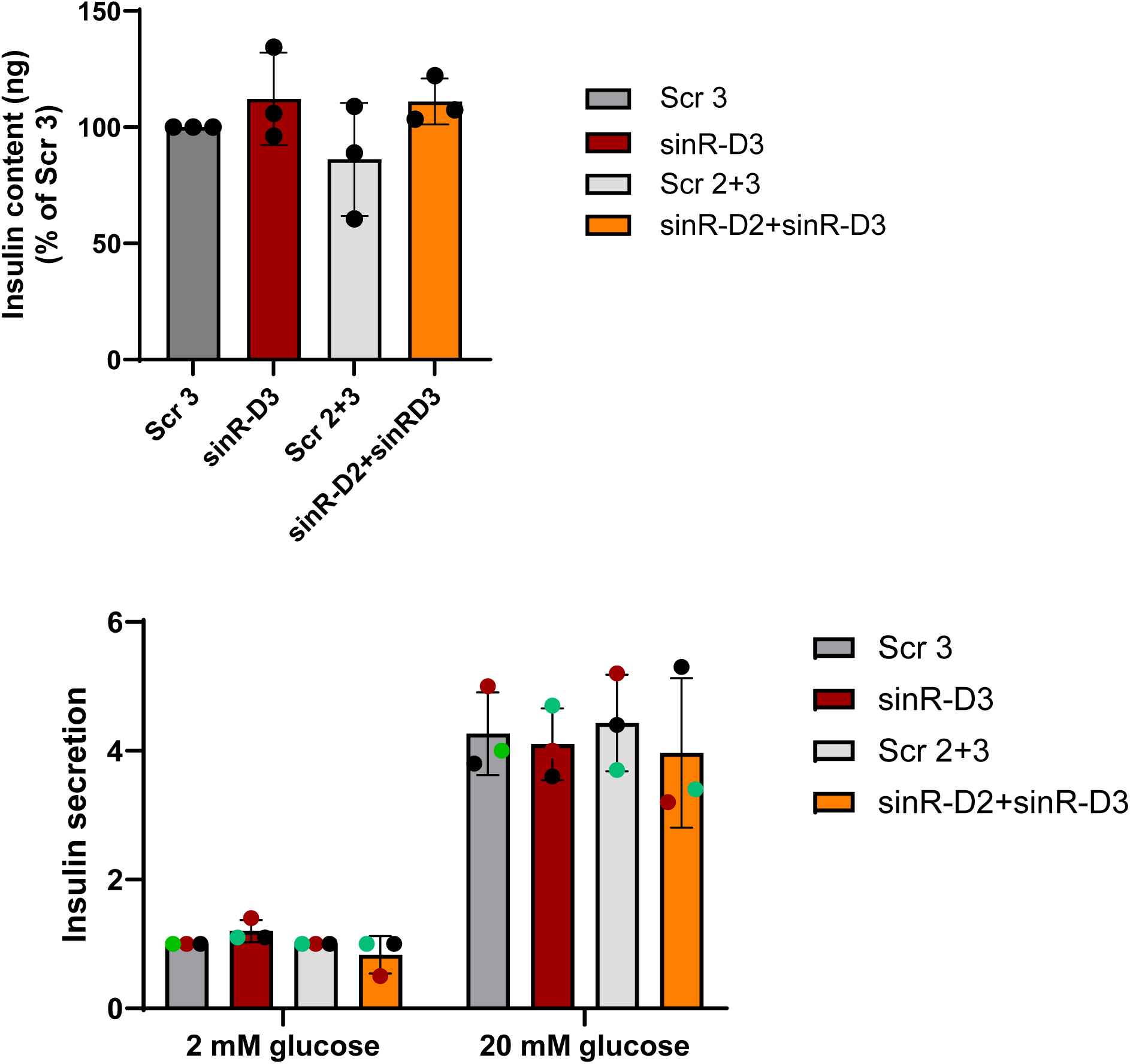
Impact of sinR-D2 and sinR-D3 on glucose-induced insulin secretion in the β-cell line MIN6. The insulin-secreting mouse MIN cell line MIN6 was transfected with sinR-D3, with both sinR-D2 and sinR-D3 or with their corresponding scrambled sequences. Insulin content (upper panel) and insulin secretion in the presence of 2 or 20 mM were measured by ELISA).

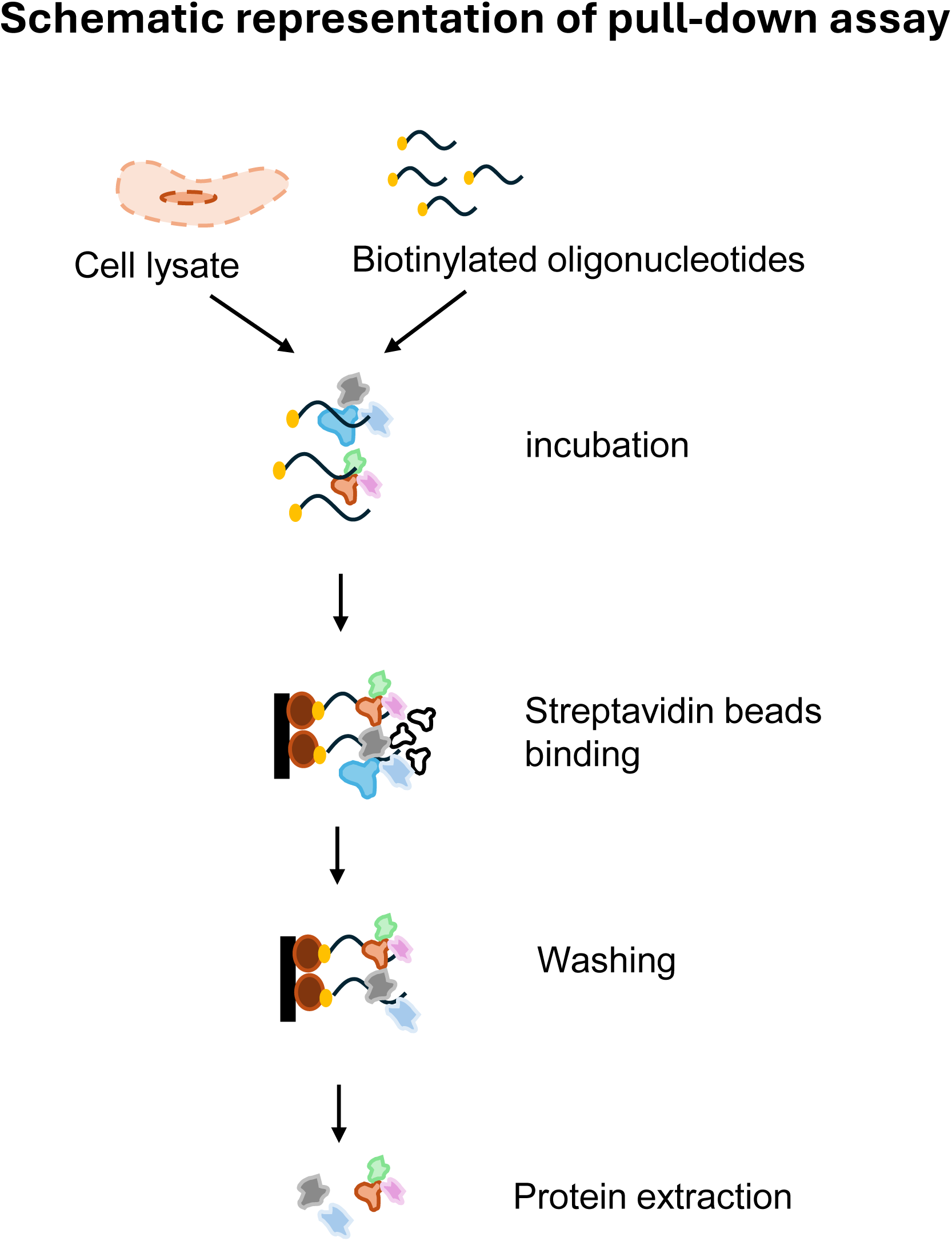

**Supplementary Table 1.**
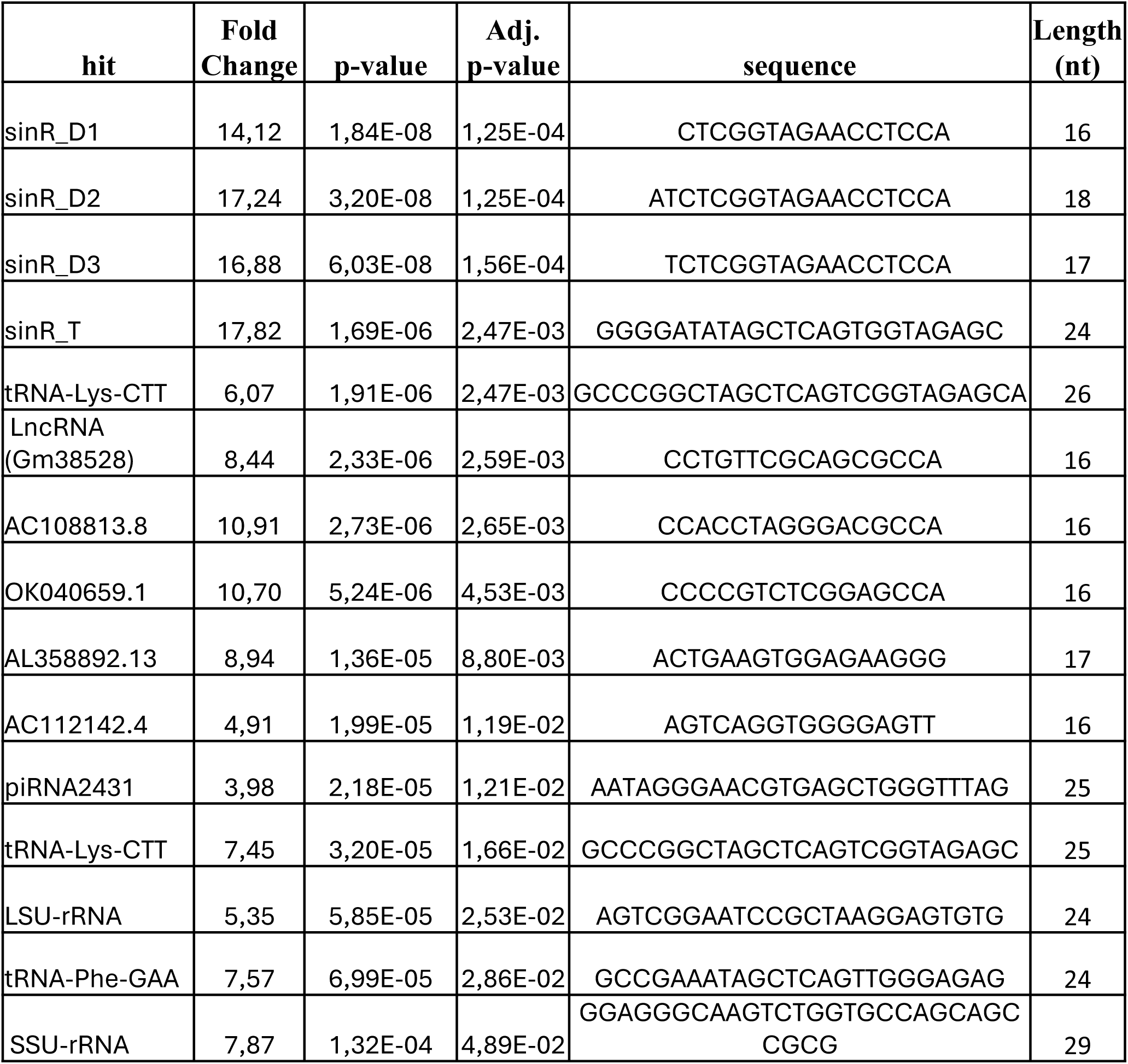
Most upregulated small ncRNA identified by RNA-seq analysis of pulled-down EU-tagged RNA in mice islets. Log_2_ fold change > 2, adjusted *p-value* < 0.05.

**Supplementary Table 2.**
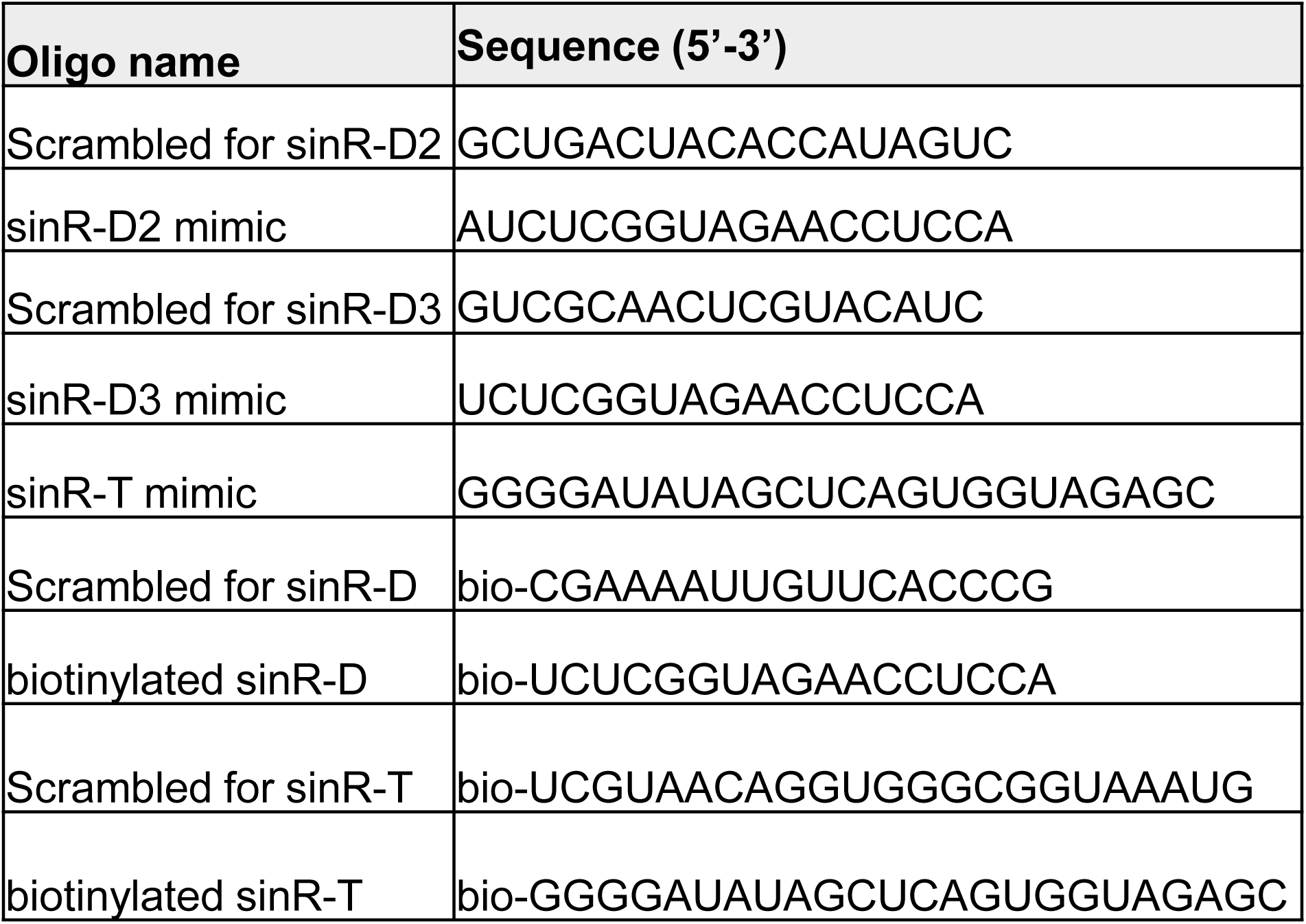
List of the oligonucleotides sequences used in the study.

**Supplementary Table 3.**
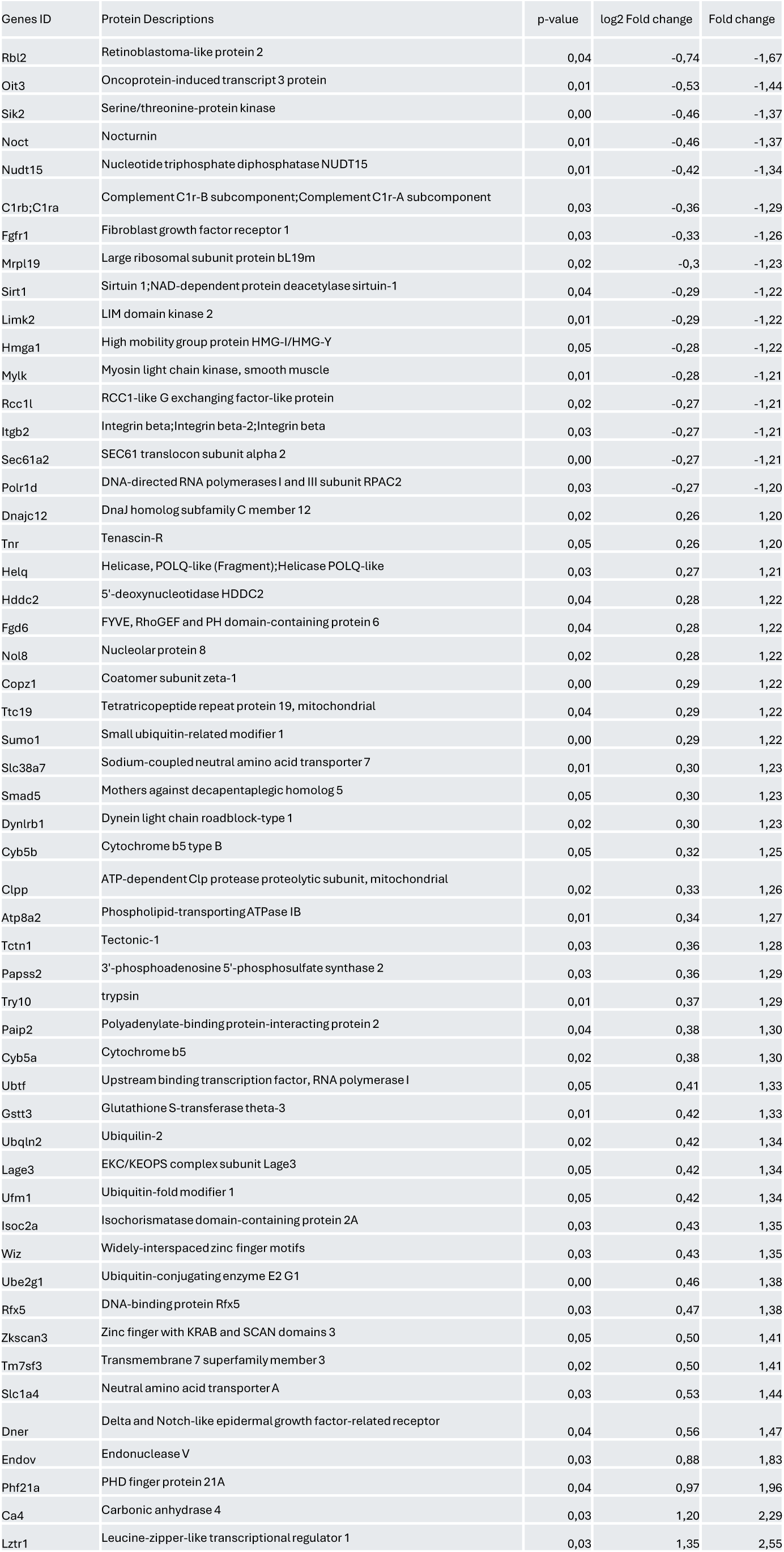
List of differentially expressed proteins identified by mass spectrometry. Fold change >1.2, *p-value* <0.05.

